# Portable framework to deploy deep learning segmentation models for medical images

**DOI:** 10.1101/2021.03.17.435903

**Authors:** Aditi Iyer, Eve Locastro, Aditya P. Apte, Harini Veeraraghavan, Joseph O. Deasy

**Author notes:** These authors contributed equally to this work (Aditi Iyer, Eve Locastro, Aditya P. Apte).

## Abstract

**Purpose:** This work presents a framework for deployment of deep learning image segmentation models for medical images across different operating systems and programming languages.

**Methods:** Computational Environment for Radiological Research (CERR) platform was extended for deploying deep learning-based segmentation models to leverage CERR’s existing functionality for radiological data import, transformation, management, and visualization. The framework is compatible with MATLAB as well as GNU Octave and Python for license-free use. Pre and post processing configurations including parameters for pre-processing images, population of channels, and post-processing segmentations was standardized using JSON format. CPU and GPU implementations of pre-trained deep learning segmentation models were packaged using Singularity containers for use in Linux and Conda environment archives for Windows, macOS and Linux operating systems. The framework accepts images in various formats including DICOM and CERR’s planC and outputs segmentation in various formats including DICOM RTSTRUCT and planC objects. The ability to access the results readily in planC format enables visualization as well as radiomics and dosimetric analysis. The framework can be readily deployed in clinical software such as MIM via their extensions.

**Results:** The open-source, GPL copyrighted framework developed in this work has been successfully used to deploy Deep Learning based segmentation models for five in-house developed and published models. These models span various treatment sites (H&N, Lung and Prostate) and modalities (CT, MR). Documentation for their usage and demo workflow is provided at https://github.com/cerr/CERR/wiki/Auto-Segmentation-models. The framework has also been used in clinical workflow for segmenting images for treatment planning and for segmenting publicly available large datasets for outcomes studies.

**Conclusions:** This work presented a comprehensive, open-source framework for deploying deep learning-based medical image segmentation models. The framework was used to translate the developed models to clinic as well as reproducible and consistent image segmentation across institutions, facilitating multi-institutional outcomes modeling studies.

## Introduction

This work fills the need for a portable software framework useful to deploy segmentation models for medical images. There are several general-purpose frameworks [PyTorch^1^, Tensorflow^2^, Caffe^3^, Keras^4^] for training deep learning-based models. There are also frameworks to deploy and distribute inference models such as DeepInfer^5^, ModelHub^6^ and NiftyNET^7^. DeepInfer, which can also be invoked from 3DSlicer^8^,currently provides segmentation models for Prostate gland, biopsy needle trajectory and tip and brain white matter hyperintensities. NiftyNET ^7^ is an open-source platform to train and deploy TensorFlow-based convolutional neural networks (CNN) models. It provides the implementation of networks such as HighRes3DNet, 3D U-net, V-net and DeepMedic. It can be used to train new models as well as share pre-trained models. While these generic frameworks offer a wide variety of network architectures, they were not designed to support specific processing requirements of medical images and porting of models between users. The framework developed in this work provides end-to-end workflow for using deep learning-based image segmentation models including conversion between data formats, visualization and simplified access to image metadata. This enables users to readily apply the trained models on datasets collected at different institutions as well as incorporate the validated models in clinical workflow.

## Methods

The framework was developed as a module within the Computational Environment for Radiological Research^9^ (CERR) software platform. This allows users to leverage existing CERR functionalities such as flexible planC data structure, visualization, feature extraction and ability to use in various programming languages including Matlab, Octave and Python via the Oct2Py bridge. Deep learning Model and framework dependencies must be consistent to ensure reproducibility of segmentation models. Container technology such as Docker and Singularity allow users to securely bundle libraries such that they are compatible with a variety of scientific computing architectures. Singularity container technology presents various advantages in terms of usage on bare-metal HPC clusters including no requirements of root privileges. While Singularity container technology completely isolates the environment form the host, it has limitations for use on Windows and Mac operating systems. Conda is a popular cross platform package and environment manager that installs and manages conda packages from the Anaconda repository as well as from the Anaconda Cloud. Conda environments can be archived and installed on other systems and locations using tools such as Conda-Pack. This is useful for deploying code in a consistent environment—potentially on systems where python and/or conda isn’t already installed. CPU and GPU implementations of pre-trained deep learning segmentation models were packaged using Singularity containers (Linux) and Conda environment archives (Windows, macOS).

**Figure 1.**
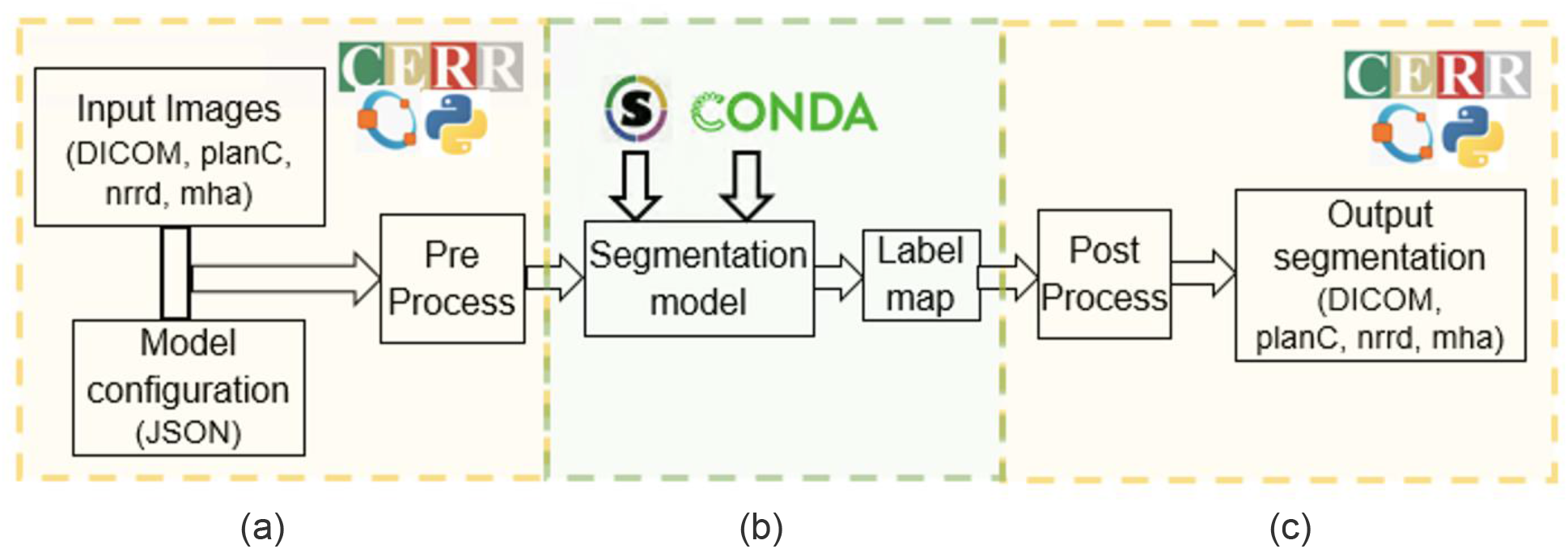
Overview of deep learning segmentation framework. (a) CERR is used in a license-free Octave or Python programming language to pre-process input images per model configuration. (b) Segmentation models are distributed as packaged Conda environments for specific operating systems (Windows, Mac, Linux) or Singularity containers for Linux. (c) Resulting segmentation labels are post-processed in CERR per model configuration.

Several pre-processing operations specific to the medical imaging domain are supported, including automated cropping to patient outline, 2D/3D resampling, 2D/2.5D/3D and multimodal channels, and orientation transformations (transverse, sagittal, coronal). The framework supports models derived with multiple image modalities. E.g. MR and CT. Various image registration options include deformable and rigid registrations as wrapper calls to popular packages such as Plastimatch^10^ and ANTS^11^. Additionally, images can be processed using Image Biomarker Standardization^12^ Initiative defined filters. Various post-processing operations are supported, including filtering to retain selected number of connected components by size or within a user-defined region of interest. Dynamic label-to-structure name maps are supported where the number output structures are variable. Table 1 lists the available processing operations. Specification of processing operations is simplified by defining them via JSON configuration file per model.

**Table 1.**
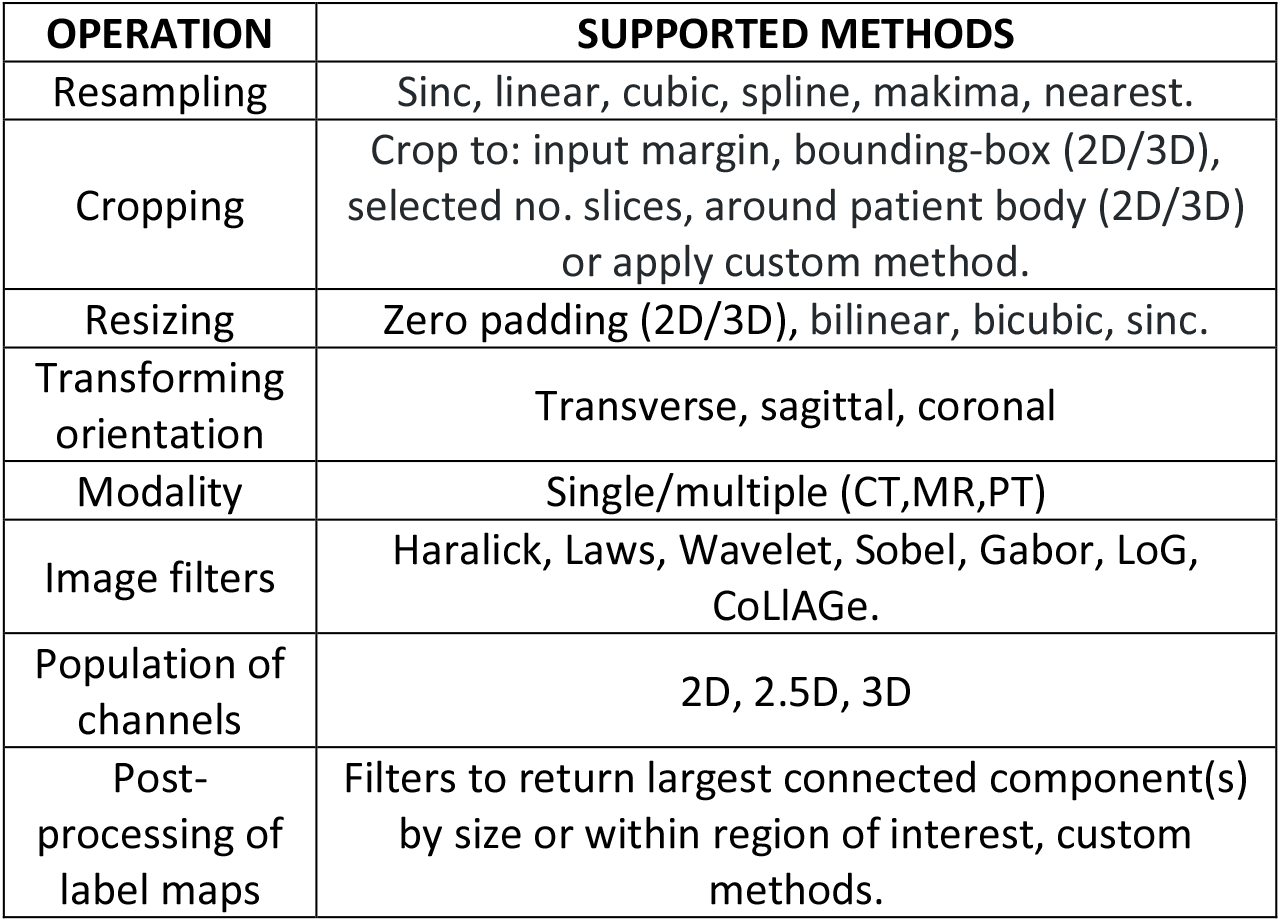
Supported pre- and post-processing methods

| OPERATION | SUPPORTED METHODS |
| --- | --- |
| Resampling | Sinc, linear, cubic, spline, makima, nearest. |
| Cropping | Crop to: input margin, bounding-box (2D/3D), selected no. slices, around patient body (2D/3D) or apply custom method. |
| Resizing | Zero padding (2D/3D), bilinear, bicubic, sinc. |
| Transforming orientation | Transverse, sagittal, coronal |
| Modality | Single/multiple (CT,MR,PT) |
| Image filters | Haralick, Laws, Wavelet, Sobel, Gabor, LoG, CoLIAGe. |
| Population of channels | 2D, 2.5D, 3D |
| Post-processing of label maps | Filters to return largest connected component(s) by size or within region of interest, custom methods. |

## Results

The segmentation framework is distributed as an open-source, GNU-copyrighted software with the CERR platform whereas the developers of models retain the copyright to their models. Documentation for available models and their usage is provided at https://github.com/cerr/CERR/wiki/Auto-Segmentation-models. A Jupyter notebook runnable on Google Colab platform demonstrates the installation of various dependencies their usage - https://github.com/cerr/CT_SwallowingAndChewing_DeepLabV3/blob/master/demo_DLseg_swallowing_and_chewing_structures.ipynb. The framework has been used to share deep learning segmentation models with several institutions within and outside the USA. Additionally, it has been used in clinical segmentation and facilitating outcomes studies^13^ using large external/open datasets.

**Table 2.** Available pre-trained models

| SITE | STRUCTURES | FRAMEWORK | MODALITY |
| --- | --- | --- | --- |
| Lung | Heart, Pericardium, Atria, Ventricles | DeepLab, Pytorch (Haq et al<sup>14</sup>) | CT |
| Prostate | Bladder, Prostate and Seminal Vesicles (CTV), Penile Bulb, Rectum, Urethra and Rectal Spacer | DeepLabV3+, Tensorflow (Elguindi et al<sup>15</sup>) | MR |
| H&N | Parotids (left, right), Submandibulars (left, right), Mandible and Brain Stem | Self-Attention, Pytorch (Jiang et al) | CT |
| Lung | Nodules | Incremental MRRN, Keras, Tensorflow (Jiang et al<sup>16</sup>) | CT |
| H&N | Masseters (left, right), medial pterygoids (left, right), larynx, and constrictor muscle | DeepLabV3+, Pytorch (Iyer et al) | CT |
| Brain | Brain metastases | Vnet, Tensorflow (Hsu et al) | CT, MR |

## Acknowledgements

This research was partially funded by NIH grant 1R01CA198121 and NIH/NCI Cancer Center Support grant P30 CA008748.

